# Federated cross-biobank conditional analysis identifies LDL-C lowering effects of *DNAJC13* haploinsufficiency and *LDLR* regulation

**DOI:** 10.64898/2026.02.03.702791

**Authors:** Harrison I.W. Wright, Liza Darrous, Lauric Ferrat, V. Kartik Chundru, Aurelie Kamoun, Andrew R. Wood, Caroline F. Wright, Kashyap A. Patel, Timothy M. Frayling, Michael N. Weedon, Robin N. Beaumont, Gareth Hawkes

## Abstract

Whole genome sequencing in diverse population-scale biobanks offers new insights into the genetic architecture of complex traits from rare and non-coding variants. However, rare single variant and aggregate associations are often confounded by linkage disequilibrium and haplotype structure, resulting in large numbers of false-positive associations. Previous methods that rely on reference panels or linkage disequilibrium-matrices to determine conditional independence in meta-analyses do not scale to very rare variants, which may be observed in only one biobank and can exhibit long-range haplotypes. Here, we implement a federated approach to perform iterative conditional meta-analysis on individual-level genotype and phenotype data across biobanks while adhering to data sharing policies. We applied our methodology to a meta-analysis of LDL-C in 614,375 individuals from UK Biobank and All of Us, encompassing six genetic ancestry groups. After conditioning, only 4.3% of significantly associated rare single variants and 6.9% of aggregates remained statistically independent. The proportion of significant aggregates that remained independent after conditioning was higher for coding-based tests than non-coding. We further validate that our approach effectively suppresses false-positive associations using simulations centred on the *LDLR* locus. We identify allelic series of variants associated with reduced LDL-C, including loss-of-function variants in *DNAJC13* and variants in the 3-prime untranslated region of *LDLR*. Our results highlight that federated conditioning can distinguish independent rare variant signals from linkage and haplotype structure artifacts in multi-ancestry meta-analyses across separate biobanks.

## Introduction

Whole genome sequences (WGS) in population settings offer a near-complete view of simple genetic variation (single nucleotide variants and short indels) across the entire allele frequency spectrum, covering both coding and non-coding regions. WGS in population-scale biobanks now cover more than one million individuals of diverse genetic ancestry, enabling multi-ancestry meta-analysis with benefits that have been extensively outlined (e.g. ^1–3^). We have previously demonstrated how aggregation of rare variants can highlight allelic series of variants in regulatory regions inaccessible by common variant genome-wide association studies (GWAS)^4,5^. We have also demonstrated that non-coding aggregate testing can highlight gene-phenotype relationships out of reach of exome sequence studies, due to differences in coverage^5^, or for genes whose exonic regions are highly constrained. In the latter case, we may only observe the association via subtler cis-regulatory effects.

However, aggregate based tests achieving study-wide significance may be false positives because of confounding due to LD and haplotype structure^5,6^. These confounding effects are particularly widespread in non-coding regions, partly because the increased scale of association testing, where an association with a phenotype may only occur via linkage or haplotype-sharing with large-effect *cis* coding variants. Previous studies, whose focus has been on common genetic variation, have used reference panels to establish statistical independence in multi-ancestry meta-analyses^1,7^. In contrast, we have previously shown that joint-modelling and stepwise conditioning of rare variants requires the WGS data from which the association statistic was derived to establish stable statistical independence^5^. Additionally, the impact of diverse-ancestry haplotype structures on rare-variant aggregate testing and conditional analysis has not been explored, particularly across multiple biobanks where we cannot aggregate individual-level genetic data. There are existing methods for cross-biobank WGS meta-analysis and common variant conditioning, which rely on aggregate or locus-level linkage disequilibrium (LD) matrices^8,9^. However, due to size constraints these are sparse matrices based on a minimum LD cutoff, which is inaccurate for rare variants^10^. Also, this approach cannot easily scale to account for long-range haplotype effects.

Our federated conditioning method solves these problems by iteratively using individual-level WGS data across biobanks to perform stepwise conditional meta-analysis (**Figure 1**). This robustly suppresses false-positive genetic associations which are driven by LD or haplotype sharing, highlighting statistically independent associations for further investigation.

**Figure 1:**
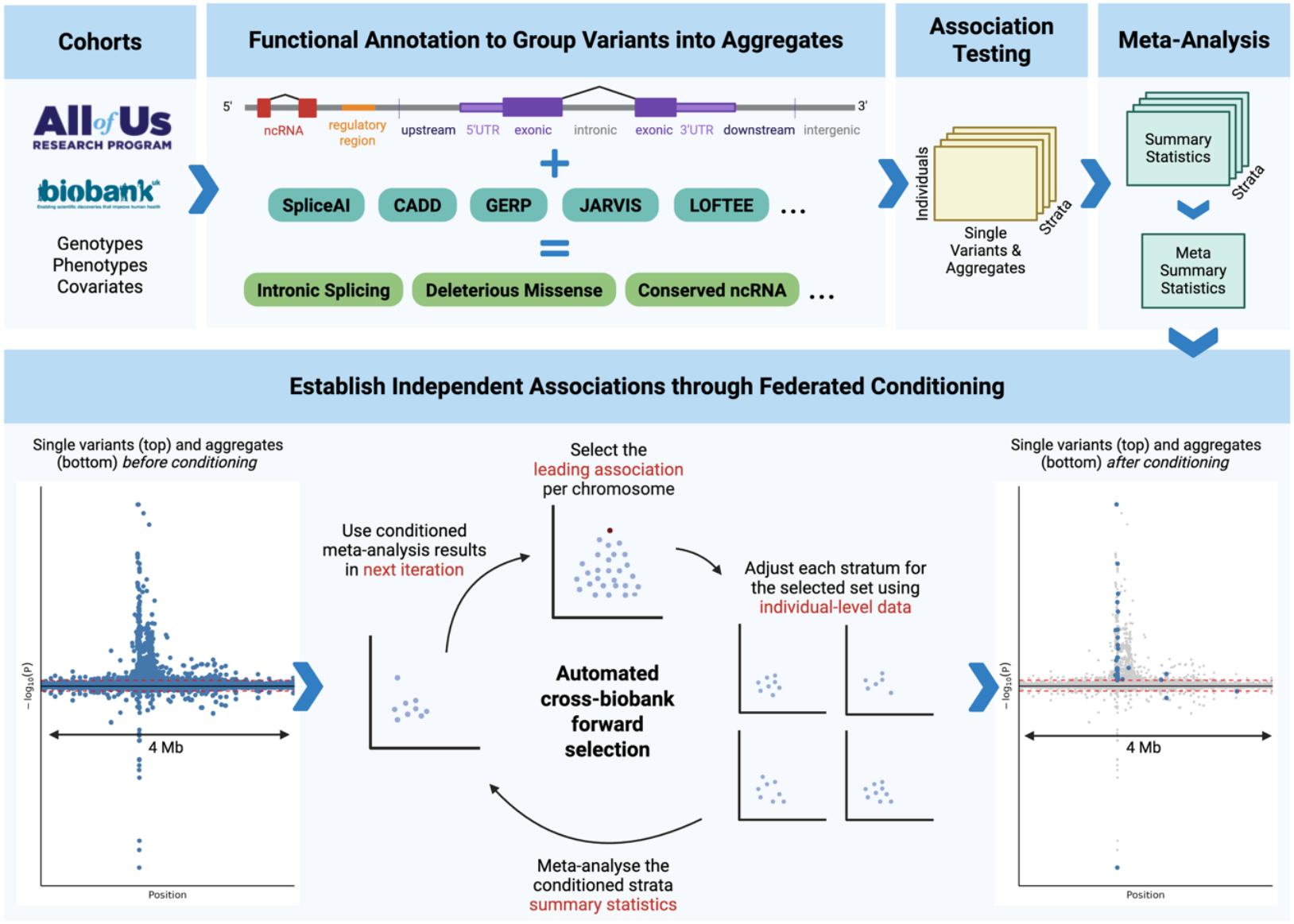
Framework for genome-wide rare variant meta-analysis and federated conditioning. Given individual-level WGS genotypes and phenotypic information (including cohort and study specific covariates), we first annotate all genetic variants according to their predicted consequences and functional annotations (see **Methods**). We perform association testing in each biobank-ancestry stratum using the respective individual-level data for both single variants and rare-variant aggregates defined by selected combinations of variant annotations. Next, we perform meta-analysis to generate baseline association statistics, where both single variant and aggregate results are confounded by linkage disequilibrium and haplotype structure as illustrated by the “before conditioning” Miami plot. We then perform forward selection using federated conditioning to establish independent associations, per chromosome (see **Methods**). Briefly, we iteratively select the top single variant or aggregate in the meta-analysis then carry out conditional analysis in each stratum, adjusting for the selected set of variants or aggregates using individual-level data, before meta-analysing at each iteration to determine the next variant or aggregate to select. We stop iterating when there are no conditionally significant associations left to select. The selected set is then taken as the set of statistically independent associations, illustrated by the blue points in the “after conditioning” Miami plot.

Low-density lipoprotein cholesterol (LDL-C) is an exemplar disease-relevant trait for which the common variant architecture has been extensively studied. Most recently, the Global Lipids Genetics Consortium conducted a meta-analysis of 1.65 million individuals for LDL-C, alongside four other blood lipid traits^1^. They identified 442 quasi-independent variants associated with LDL-C and demonstrated that common variant polygenic risk scores can explain up to 16% of the variance in LDL-C. Interpreting the biological mechanisms underlying many of these common variant loci is difficult because they often lie in non-coding regions of the genome, with no clear effector gene or mechanism. We hypothesised that WGS analysis including rare variants in non-coding regions will help elucidate the pathways by which these LDL-C loci act.

Here, we perform meta-analysis and federated conditioning of rare single variants, as well as rare coding and non-coding variant aggregates, for LDL-C across UK Biobank^11,12^ (N = 462,094) and All of Us^13,14^ (N = 152,281), covering six genetic ancestry groups: South Asian (SAS; N_UKB = 9,621; N_AOU = 1,790), African (AFR; N_UKB = 7,388, N_AOU = 28,552), Admixed American (AMR; N_AOU = 21,512 ), East Asian (EAS; N_AOU = 3,200), Middle Eastern (MID; N_AOU = 577) and European (EUR; N_UKB = 445,085; N_AOU = 96,650). By analysing the whole genome and accounting for haplotype and linkage effects, we further our understanding of the genetic aetiology of LDL-C.

## Results

### A framework for rare variant meta-analysis and federated conditioning

We developed a meta-analysis and federated conditioning framework (**Figure 1**) to address several methodological challenges with genome-wide rare variant association testing, including the large number of variants tested (204.9 million for LDL-C), reduced power compared to common variants, false-positives caused by long-range LD and haplotypes, and multiple independent associations hidden in a large association peak.

To handle the scale of the analysis, we use REGENIE^15^ in parallel across chunks of the genome, with number of variants per chunk optimised based on stratum size. We perform single-variant association tests with a minor-allele-count (MAC) of 11 or more in at least one stratum, as well as rare-variant aggregate burden tests (minor allele frequency; MAF < 0.1%). To determine which variants to group together for aggregate testing, we use combinations of positional and functional and annotations across both coding and non-coding regions (**Methods**).

Additive burden tests are most well-powered when all variants act homogeneously on the phenotype, for example in a coding loss-of-function aggregate, while aggregated Cauchy association (ACAT) tests^16^ are better powered when variants act bi-directionally and sparsely, for example in a regulatory aggregate where not all annotated variants are causal. Using both types of test increases discovery across the coding and non-coding genome. In our framework, we perform fixed-effects meta-analysis of single variant and rare-variant burden tests using GWAMA^17^ and generate ACAT aggregate summary statistics by combining the meta-analysed single variants (MAF < 0.1%) and an additional MAC < 11 burden, using GCTA^18^.

To establish independent associations and suppress false-positives we developed workflows which automate forward selection of meta-analysis results using federated conditioning (**Methods**). Briefly, per-chromosome at each iteration of conditioning, we select the top variant/aggregate in the meta-analysis by p-value. Then, using REGENIE and individual-level data, we re-analyse unselected variants/aggregates (limited to those passing a sub-significant p-value threshold) in each stratum separately, adjusting for the selected set. We then meta-analyse the adjusted summary statistics and add the next variant/aggregate to the selected set, repeating the process until no additional conditionally significant associations remain. For single variants, at each iteration, we additionally prune based on per-stratum collinearity with already-selected variants (r^2^ < 0.9), in line with GCTA-CoJo^19^.

### Federated conditioning identified 120 rare single variants independently associated with LDL-C levels

We aimed to determine the added value of our multi-ancestry federated conditioning approach for identifying rare single variant associations. Our meta-analysis of 614,375 people highlighted 74,514 genetic variants associated with LDL-C (*P* < 2.95e-10), of which 2,821 (3.79%) were rare (MAF < 1%). After conditioning (**Methods**), we identified 559 independent single variants associated with LDL-C, of which 120 (21.5%) were rare (**Figure 2a; Supplementary Table 1**). These included 79 rare coding variants, highlighting 39 genes, and 41 rare non-coding variants. Concordance of effect sizes between strata are provided in **Supplementary Figure 1**.

**Figure 2:**
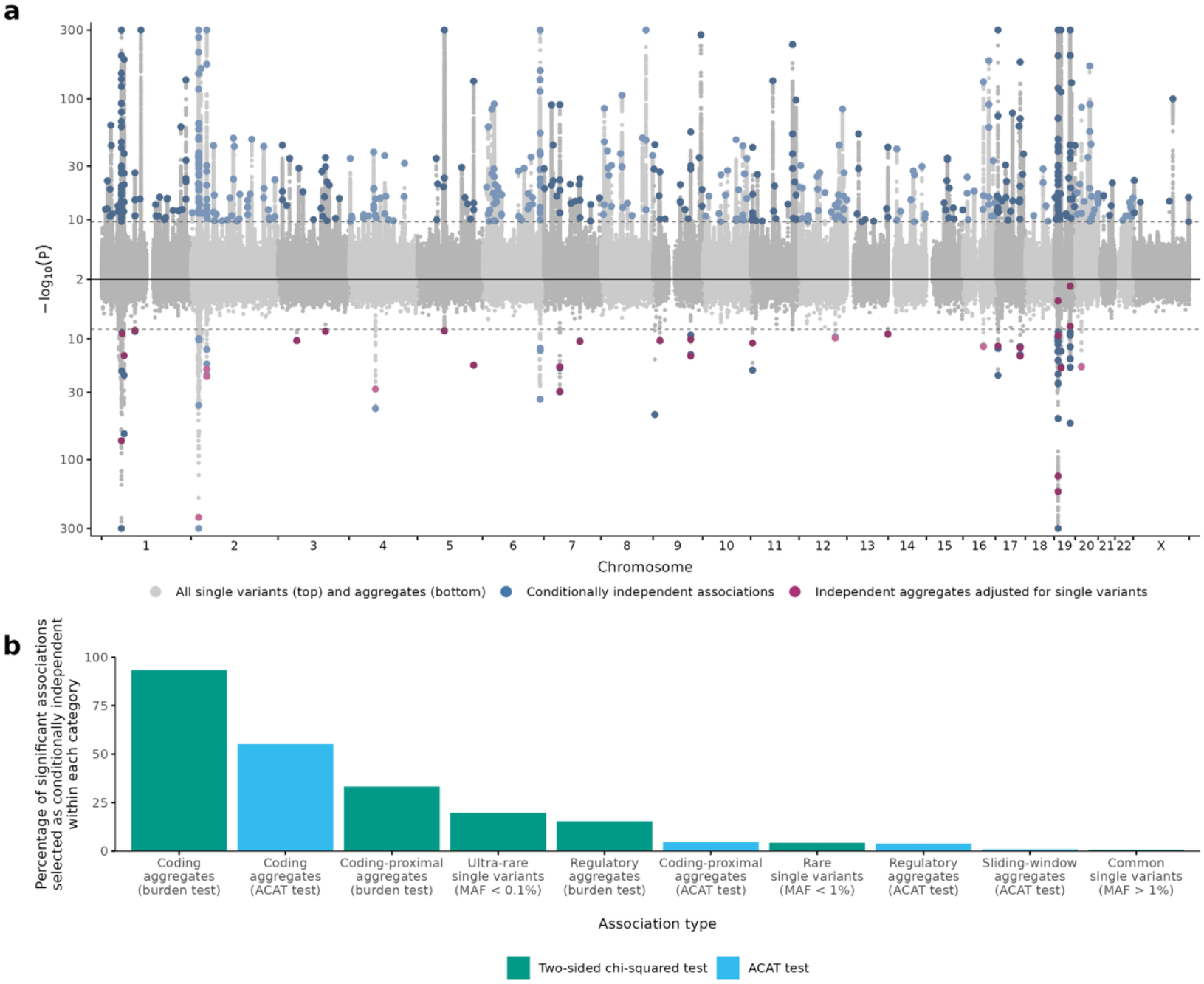
Federated conditioning supresses non-causal genetic associations with LDL-C. **a)** Miami plot of single variants (positive y-axis) and rare-variant aggregates adjusted for previously known common variants (negative y-axis), with y-axis magnitude representing -log_10_(*P*) of association. The x-axis represents chromosomal position in sequential chromosomes. P-values are based on a two-sided chi-squared test or ACAT test and are restricted to 1e-300 < *P* <1e-2 for display. Dashed lines represent study-wide significance for each analysis. Grey points represent genetic associations which were not selected as independent and blue points represent conditionally independent associations. Purple points represent blue independent aggregate associations after additionally adjusting for independent single variant associations. **b)** Bar chart showing the percentage of significant unconditioned associations which were selected as independent by our federated conditioning approach, split by frequency for single variants and category/test type for aggregates. Overall, rare variants and coding aggregates were more likely to be conditionally independent.

Rarer variants were more likely to be statistically independent than common variants (**Figure 2b; Supplementary Figure 2**). Overall, 439/71,693 (0.6%) of common variants, 120/2821 (4.3%) of rare variants and 36/184 (19.6%) of MAF < 0.01% variants remained significant after our conditional analysis. Rare variants were also more likely to be independent of previous GWAS hits. After adjusting for variants previously reported to be associated with LDL-C^1^, 182/439 (41.5%) of common and 107/120 (89.2%) of rare independent variants remained significant.

To ensure statistical independence of our single variant associations, in light of known issues with forward selection^20^, we performed additional joint-modelling of the genetic variants selected by federated conditioning, using GCTA-CoJo in each stratum, and meta-analysed the joint-statistics. Our results confirmed a high degree of independence between selected variants: the maximum joint p-value was 2.78e-07 and 534/559 (95.5%) of variants remained study-wide significant, of which 105 were rare.

For example, we identified a rare highly-deleterious loss-of-function variant in *RORC* which reduces LDL-C (1:151831737:G:A; beta = -0.100 SD; se = 0.0148; *P*_*indep*_ = 1.37e-11, freq = 0.0031, CADD = 36.0). However, an aggregate of loss-of-function variants in *RORC* did not associate with LDL-C (*P* = 0.374). The variant also overlaps three non-MANE-select *RORC* transcripts: the 5’UTR of ENST00000652040.1, and the non-protein-coding transcripts ENST00000638901.1 and ENST00000651814.1. This suggests it could be acting through mechanisms other than loss-of-function. We additionally identified rare large-effect variants enriched in specific ancestries, including an 8-basepair insertion in *SERPINB11*, only common enough to test within the AMR genetic ancestry group (18:63712688:C:CATCAGGTA; beta = -0.411 SD; se = 0.0680; *P*_*indep*_ = 2.21e-10; freq = 0.0041).

### Most statistically significant aggregates are driven by LD and haplotype structure

We next applied our conditioning framework to rare-variant aggregate testing. In our initial meta-analysis, 142 rare-variant burden and 1,803 rare-variant ACAT associations achieved study-wide significance (*P* < 8.71e-9) after adjustment for previous GWAS variants. After conditioning, only 29/142 (20.4%) of burden and 32/1803 (1.8%) of ACAT associations remained significant (**Figure 2a; Supplementary Table 2**).

Non-coding aggregates were much more likely to be supressed by conditioning than coding aggregates. To quantify this, we first merged overlapping aggregate definitions into four classifications: coding; proximal (to coding); regulatory; and sliding windows which exclude coding and proximal variants (**Figure 2b; Supplementary Table 3**). For burden tests, 15.4% of regulatory and 33.3% of proximal aggregates were independently significant, compared to 93.3% of coding aggregates. For ACAT tests, only 3.8% of regulatory, 4.6% of proximal and 55.2% of coding aggregates were independent. Sliding windows were only associated through ACAT tests, with just 0.9% being independent of other results. Overall, 6.9% of associations were selected as independent by conditioning. Non-coding aggregates are less likely to be independent because the large-effect casual aggregates which cause correlated aggregates to be spuriously associated are often coding. ACAT tests are less likely to be independent than burden tests, likely because ACAT tests are more sensitive to individually highly associated single variants.

To further evidence the confounding effect of LD and haplotype structure, we performed a simulation study. We generated phenotypes which were causally associated exclusively with coding variation in *LDLR* (low density lipoprotein receptor), with varying effect size distributions, and performed association testing for all aggregates on chromosome 19 (**Methods**). We found that non-causal aggregates over 2 Mb away associated with our simulated phenotypes (**Supplementary Figure 3a**). Our federated conditioning approach supressed these spurious associations (**Supplementary Figure 3b**).

**Figure 3:**
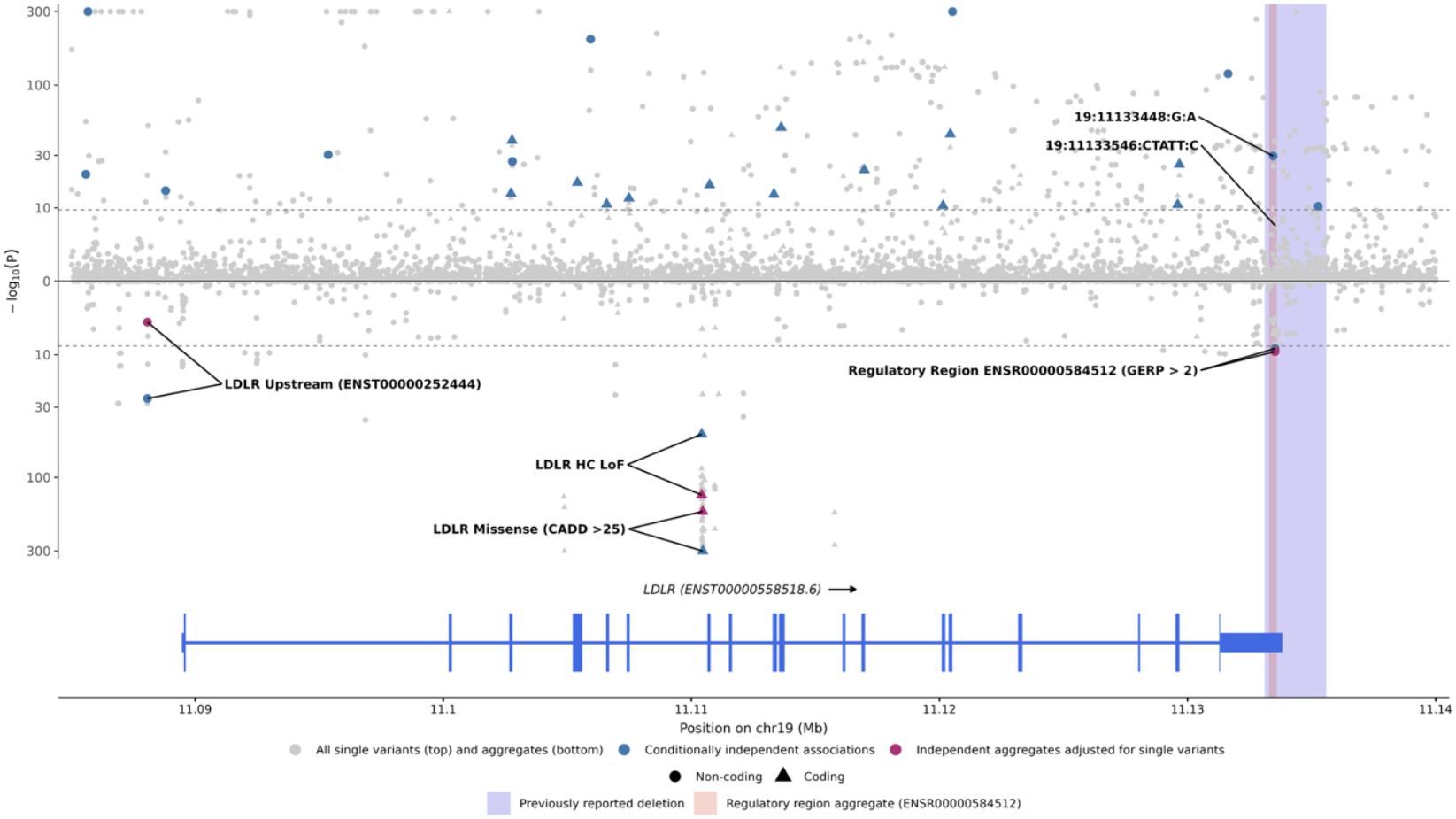
Summary of association between rare 3’UTR variants in *LDLR* and LDL-C. Miami plot of single variants (positive y-axis) and rare-variant aggregates adjusted for previously known common variants (negative y-axis), with y-axis magnitude representing -log_10_(P) of association. P-values are based on a two-sided chi-squared test or ACAT test and are restricted to *P* > 1e-300 for display. Dashed lines represent study-wide significance for each analysis. Grey points represent genetic associations which were not selected as independent and blue points represent conditionally independent associations. Purple points represent blue independent aggregate associations after additionally adjusting for independent single variant associations. Triangular points represent coding associations (within an exon of the *LDLR* transcript shown) and circular points represent non-coding associations. The shaded red region shows the extent of Ensembl (v110) predicted regulatory region ENSR00000584512, and the shaded blue region represents a previously reported 2.5 kb deletion.

We also prioritised aggregate associations with LDL-C by adjusting for our independent single variants, which included the *APOE* e2/e3/e4 haplotypes defined by allelic combinations of 19:44908822:C:T and 19:44908684:T:C. This left 31 aggregate associations that were most likely to highlight causal effects not previous identified by GWAS or our single variant meta-analyses (**Supplementary Table 2**).

### An aggregate of loss-of-function variants in *DNAJC13* reduces LDL-C across ancestries, fine-mapping common variant associations

We identified an association between loss-of-functions variants in *DNAJC13* (DnaJ heat shock protein family (Hsp40) member C13) and reduced LDL-C (beta = -0.396 SD; se= 0.0671; *P*_*adjCommon*_ = 3.90e-9; freq = 1.53e-4), replicating across the four genetic ancestries where it was common enough to test (**Supplementary Table 4**). *DNAJC13* is a highly constrained gene (pLI = 1; o/e=0.21 [0.17-0.27]) with 56 exons and improved coverage in whole-genome sequencing compared to exome sequencing (**Supplementary Figure 4**). Multiple common genetic variants in the intronic regions of *DNAJC13* have previously been associated with LDL-C and total cholesterol in diverse samples, and coding variants have been linked with Parkinsons disease^21^. *DNAJC13* is widely expressed, including in haemoglobin, and has been linked to endosomal trafficking^22^, suggesting it could alter lipid trafficking between cells.

**Figure 4:**
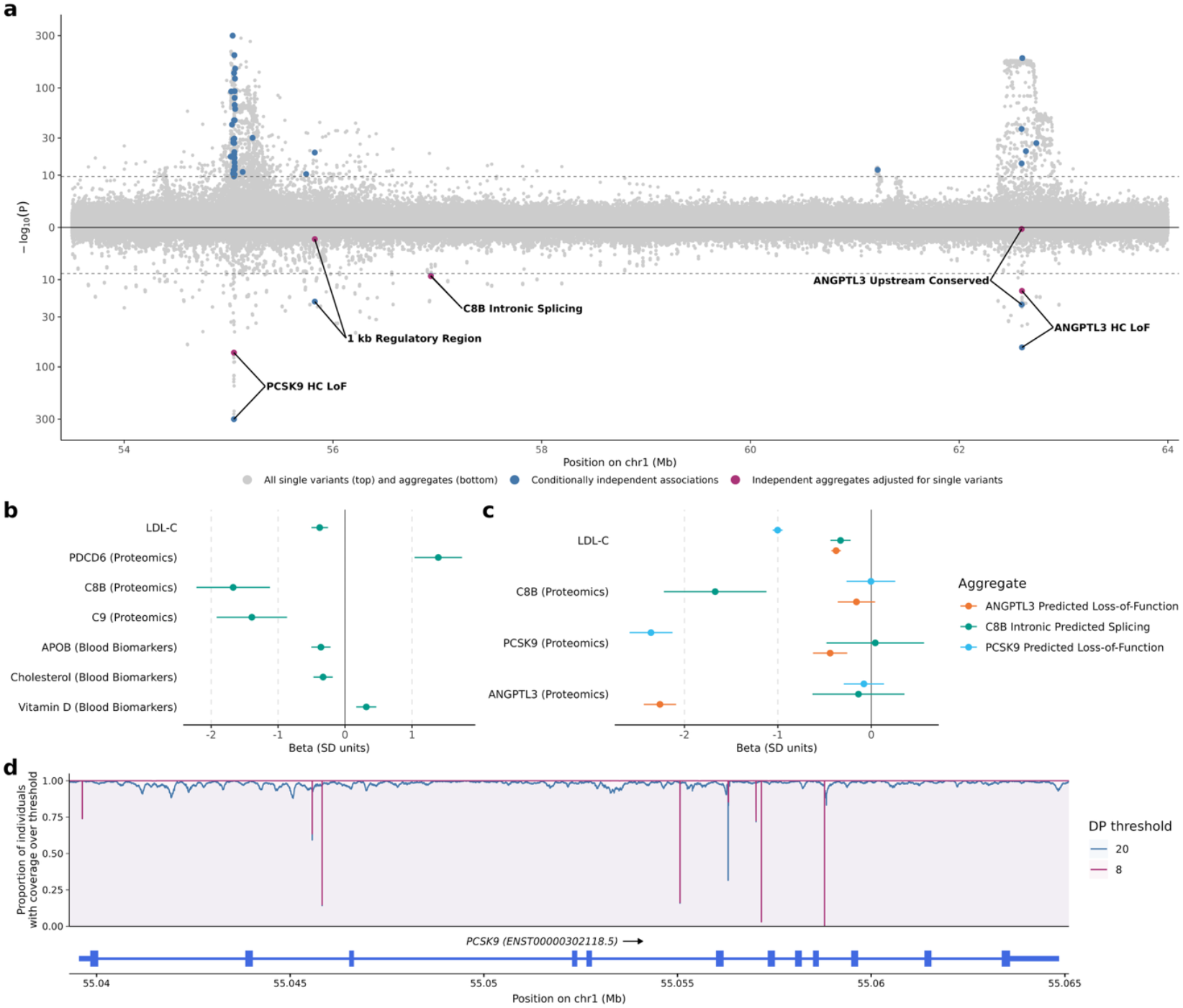
Summary of *C8B* locus and its haplotype links to *PCSK9*. **a)** Miami plot of single variants (positive y-axis) and rare-variant aggregates adjusted for previously known common variants (negative y-axis), with y-axis magnitude representing –log_10_(P) of association. P-values are based on a two-sided chi-squared test or ACAT test and are restricted to *P* > 1e-300 for display. Dashed lines represent study-wide significance for each analysis. Grey points represent genetic associations which were not selected as independent and blue points represent conditionally independent associations. Purple points represent blue independent aggregate associations after additionally adjusting for independent single variant associations. **b)** Forest plot showing significant associations from a phenome-wide scan of the *C8B* intronic splicing rare variant aggregate **c)** Forest plot showing associations of the independent rare variant aggregates in the *PCSK9-C8B-ANGPTL3* locus with LDL-C and the three gene’s protein levels. **d)** UKB WGS coverage for *PCSK9*, showing intronic gaps in coverage where a low proportion of individuals have sequncing depth (DP) above the thresholds of 20 (blue) and 8 (purple).

### Rare regulatory variants in the 3’UTR of *LDLR* reduce LDL-C

We identified an association between reduced LDL-C and an aggregate of highly-conserved (GERP > 2) variants in a regulatory region overlapping the 3’UTR of *LDLR* (ENSR00000584512; 19:11,133,281-11,133,600; beta = -0.866 SD; se = 0.144; *P*_*adjCommon*_ = 1.74e-09; freq = 3.7e-5), as well as a rare single variant in the same region (19:11133448:G:A; beta = -0.200 SD; se = 0.0175; *P*_*indep*_ = 2.06E-30; freq = 2.23e-3). These non-coding variants contrast with *LDLR* loss-of-function (beta = 1.41 SD) and missense (beta = 0.940 SD) aggregates, which independently associate with increased LDL-C. A high constraint score (JARVIS = 0.995; UKB Depletion Score = 0.976) four base-pair rare deletion was responsible for a significant portion of the regulatory aggregate signal (19:11133546:CTATT:C; beta = -1.087 SD; se = 0.186; *P*_*indep*_ = 5.81e-9; freq = 2.6e-5). Previously, a 2.5 kb LDL-C-lowering deletion, also overlapping the 3’UTR of *LDLR*, was observed in 7 heterogenous carriers from the same family^23^. Disruption of microRNA binding sites was suggested as the key mechanism by which this deletion reduced LDL-C. The variants identified here are likely to act through the same mechanism, but affect fewer, more specific sites, fine-mapping the association. Other more common variants which alter *LDLR* splicing have also been shown to reduce LDL-C^24^, suggesting that carefully targeted regulation of *LDLR* could reduce cholesterol levels.

### Rare intronic predicted splicing variants in *C8B* are associated with reduced LDL-C, and reveal challenges in aggregate interpretation

We also identified an association between predicted intronic splice variants (SpliceAI > 0.5) in *C8B* (complement C8 beta chain) and reduced LDL-C (beta = -0.381 SD; se = 0.0629; *P*_*adjCommon*_ = 1.40e-9) (**Figure 4a**). *C8B* shows no evidence of constraint in GNOMAD^25^ (pLI = 0), and we observed no evidence of association for missense (*P*_*Burden*_ = 0.0811; *P*_ACAT_ = 0.033) and loss-of-function variants (*P*_*Burden*_ = 0.467; *P*_ACAT_ = 0.0827) in *C8B*.

*C8B* is almost exclusively expressed in liver tissue^26,27^ and, in a phenome-wide scan within UKB (**Figure 4b; Supplementary Table 5**), these predicted intronic splice variants associated with circulating protein levels of *C8B* (beta = -1.67SD; *P* = 2.37e-9), independently of coding variants in *C8B*^*5*^. The aggregate also associated with circulating levels of C9 (beta = -1.39SD; *P* = 2.15e-7) and PDCD6 (beta = 1.39; *P* = 1.33e-14). A single variant contributing to the burden, 1:56943691:C:G, previously linked to decreased metabolomic Apolipoprotein B levels^28^, is replicated here (beta = -0.360 SD for LDL-C; *P* = 9.23e-7).

However, we observed evidence of a putative long-range haplotype effect between *C8B* and *PCSK9* (**Supplementary Table 6**). Although the r^2^ correlations between the *C8B* aggregate and coding *PCSK9* variants were low, D’, a measure of haplotype sharing, shows significant correlation between the *C8B* intronic splice variants and several coding *PCSK9* variants 1.9 MB away. Our federated conditioning approach aims to handle these haplotype effects and shows that the *C8B* aggregate association remains significant after adjusting for the *PCSK9* loss-of-function aggregate. Additionally, we found no evidence of association between rare intronic *C8B* variants and levels of circulating *PCSK9* or *ANGPTL3* in UKB (N = 40,130; P > 0.116; **Figure 4c**). However, there could be a small-effect association we do not have the power to detect. Also, our conditioning approach could be affected by gaps in coverage within the introns of *PCSK9* (**Figure 4d**), which could be masking causal variants. It is not possible to robustly condition on missing data, so the possibility remains that the *C8B* aggregate could be driven by a long-range haplotype effect with *PCSK9*, although untagged causal variants would still have been expected to have impacted measured circulating protein levels if they shared a haplotype.

## Discussion

We have developed a framework for both single variant and aggregate meta-analysis and forward selection by federated conditional analysis across biobanks, with WGS data. To determine the statistical independence of very rare single variants it is not feasible to use methods such as GCTA-CoJo, which rely on a reference panel. To condition rare-variant aggregates it is not practical to employ methods that use LD matrices, which do not scale to long range rare-variant haplotypes and break down for ultra-rare variants. We show that a powerful alternative for rare single variants is an iterative approach where conditional analyses are run separately in each cohort and meta-analysed, top variants selected, and further analyses run until there are no significant variants remaining to select. We have automated this process so that hundreds of iterative steps can be run across UKB and AoU with a single command-line argument. We take the same approach for aggregate conditioning, building on previous work where we conditioned aggregates by forward selection in a single cohort. This framework can be easily extended to other biobanks that provide or allow API access to upload inputs, launch analysis workflows and download summary statistics.

We show that, as with common single variant loci, most *cis* aggregate associations are driven by LD and haplotype structure, requiring conditional analysis to establish independence. In simulations, a single causal aggregate at the *LDLR* locus caused aggregates to be associated over 2 Mb away, with conditioning supressing these associations. For LDL-C, conditioning reduced the total number of rare single variant and rare-variant aggregate associations by 95.7% and 93.1%, respectively. We show that interpretation of non-coding aggregates requires particular caution, with only 4.6% of coding-proximal ACAT associations being independent. In contrast, 28 out of 30 coding aggregate burden tests were independent, but this still represents a false-positive rate of 6.7%. For example, a coding association in *ABCG8* was driven by coding variants in *ABCG5*.

We identified several rare variants and aggregates that independently lower LDL-C, including loss of function variants in *DNAJC13* which fine-map a GWAS locus and regulatory variants in the 3’ UTR of *LDLR* which may be acting through disruption of microRNA binding sites. Rarer variants tend to have larger effects, and fewer LD-partners, making them good targets for further study of the underlying biological mechanism and drug development. Additionally, conditioning on previous GWAS variants prioritised novel, independent rare single variants and aggregates for further study.

There are several limitations in our study. Firstly, UKB and AoU are both population cohorts with a healthy volunteer recruitment bias and may be depleted of individuals with extreme LDL-C levels. The LDL-C phenotype in AoU may also be biased due to being derived from electronic health records, whereas UKB LDL-C was measured at baseline. Secondly, our federated conditioning is a forward selection procedure, which may not select all causal variants and aggregates. It may also bias regression coefficients, especially in the presence of collinearity^20^. However, we attempt to mitigate this issue by only selecting variants that are not collinear with already selected variants in single variant conditioning, by dropping variants and aggregates that fall below a significance threshold from further consideration, and by taking advantage of REGENIE’s computation of an orthonormal transformation of the covariates (including conditional variants), which is projected out of the individual-level genotypes^15^. Additionally, the main aim of the conditioning is not to develop a regression model for prediction, rather it is to select associations for further study, where we check the conditioned p-values and betas against the unconditioned, and perform further downstream analyses. Thirdly, not all regions of the genome are well covered, and only single nucleotide variants and short indels are called in the short-read WGS data used in this study. As a result, some significantly associated single variants and aggregates may be tagging causal variants that are not present in the data. Finally, 1% of samples in the v8 release of AoU did not pass the ≥ 30x WGS mean coverage threshold but were mistakenly included in the release. This issue was identified by AoU after we had run our analyses, so these samples are included in this study.

In summary, we show that without conditioning, many significant rare variant associations in large-scale WGS association analyses are driven by LD and haplotype structure. We developed a federated conditioning framework which prioritises independent single variants and aggregates. Using this framework, we identified strong LDL-C lowering effects, providing good targets for further study of the underlying biological mechanisms and drug development.

### Acknowledgements, Funding, Author Contributions

HW and MNW are supported by Medical Research Council grant MR/Y003748/1. ARW is supported by the Academy of Medical Sciences / the Wellcome Trust / the Government Department of Business, Energy and Industrial Strategy / the British Heart Foundation / Diabetes UK Springboard Award [SBF006\1134]. TMF is supported by MRC awards MR/WO14548/1 and MR/T002239/1. GH is supported by the Medical Research Council grant UKRI327. The research utilised data from the UK Biobank resource carried out under UK Biobank application number 103356. UK Biobank protocols were approved by the National Research Ethics Service Committee. The authors would like to acknowledge the use of the University of Exeter High-Performance Computing (HPC) facility in carrying out this work, funded by an MRC Clinical Research Infrastructure award (MRC Grant: MR/M008924/1). This study was supported by the National Institute for Health and Care Research Exeter Biomedical Research Centre. The views expressed are those of the authors and not necessarily those of the NIHR or the Department of Health and Social Care. We gratefully acknowledge All of Us participants for their contributions, without whom this research would not have been possible. We also thank the National Institutes of Health’s All of Us Research Program for making available the participant data examined in this study.

## Data Availability

Individual-level data cannot be shared publicly because of data availability and data return policies of the UK Biobank and All of Us. Data are available from the UK Biobank and All of Us for researchers who meet the criteria for access (http://www.ukbiobank.ac.uk; https://allofus.nih.gov/). Summary statistics are available at [TO BE ADDED UPON ACCEPTANCE].

## Code Availability

Code used for analysis and figure generation is available on GitHub [https://github.com/ExeterGenetics/federated-conditioning-ldl-c-2026] (Currently the looper module is available publicly, full code will be uploaded and zenodo snapshot created upon acceptance).

## Methods

This research complies with all appropriate ethical regulations. Ethics approval for the UK Biobank study was obtained from the North West Centre for Research Ethics Committee (11/NW/0382).

### UK Biobank and All of Us Whole Genome Sequencing

The whole genome sequencing performed for UKB had an average coverage of 32.5X using Illumina NovaSeq 6000 sequencing machines^12^. The genome build used for sequencing was GRCh38: single variant nucleotide polymorphisms and short ‘indels’ were jointly called using DRAGEN 3.7.8.

We set any UKB-WGS genotype calls to missing if either the sum(LAD) < 8 (local allele depth; LAD) per sample-genotype or GQ < 10 (genotype quality; GQ) for each of the 154,430 pVCFs provided by UKB using bcftools^29^. After these additional quality control steps, the transmission rate of singletons, which should theoretically be exactly 0.5 (assuming the majority of variants are not under strong negative selection), was 0.497, as compared to 0.456 as originally provided by UKB.

The whole genome sequencing performed for AoU had average coverage 37.9X using Illumina NovaSeq 6000 sequencing machines. The genome build used for sequencing was GRCh38: single variant nucleotide polymorphisms and short ‘indels’ were called using DRAGEN 3.7.8, followed by joint calling using the Genomic Variant Store^13^.

We exported the All of US WGS data from the provided VDS format to VCF format using HAIL^30^ v0.2.126. We subsequently removed variants where ‘FILTER!=PASS’ (AoU site level filtering^31^; FILTER) and set genotypes to missing where ‘FT==FAIL’ (AoU VETS^32^ filtering; FT) using bcftools^29^.

For both All of Us and UK Biobank, we subsequently dropped any variant with a missingness greater than 10%. Indels were normalised and left-aligned using bcftools based on a 1000G b38 reference available at https://ftp.1000genomes.ebi.ac.uk/vol1/ftp/technical/reference/GRCh38_reference_genome/ (accessed 13/02/2025). Finally, a multi-allele splitting procedure was applied, and each variant was assigned a unique ID (CHR:BP:REF:ALT) before conversion to PLINK^33^ (v2.0.0-a.6.9) p(gen/var/sam) format and merging of all shards per chromosome.

### LDL-C Definition in UK Biobank

We defined LDL based on UKB field 30780 instance 0; for those with a missing value we used instance 1. To adjust for statin and cholesterol medication, we extracted fields 6177, 6153 and 20003. We inflated an individual’s measured LDL by a factor of 1.4068 if they were on statins, defined as the participant having records, or if they had a record of having been prescribed ezetimibe.

### LDL-C Definition in All of Us

Extracting phenotype data for LDL-C consisted of an initial sample filtering step where we removed: 1) 987 (0.2%) samples flagged as population outliers based on sequencing metrics, as described by the All of Us Genomic Quality Report v8^31^, 2) one sample from each pair of monozygotic twins (or sample duplicates), defined as having kinship > 0.46 in the Hail pc_relate output provided by AoU. We did not remove 4,044 samples which failed AoU’s ≥ 30x coverage threshold because this issue was identified by AoU after we had run our analyses, with 1,373 of these samples included in our final LDL-C phenotype.

After extracting all lipid measurements from Electronic Health Record (EHR) data for the remaining samples, we removed measurements where the participant was younger than 18 or older than 75 years of age at the time of measurement^34^.

For the remaining lipid measurements, we converted their units to mmol/L and filtered out any measurements equal to or below 0 mmol/L (N =1449), and further any outlier measurement (LDL-C greater than 27.75 mmol/L) using the known cutoff from the Vanderbilt University Medical Center EHR study^34^, leaving us with 1,006,348 LDL-C measurements.

We next aimed to adjust lipid measurements that fell within a lipid-lowering drug (LLD) intake time frame. In order to do so, we categorized our measurements into three groups as follows: 1) measurements of samples who do not have a record of taking LLD (no adjustment), 2) measurements of samples who have records both while taking or not taking LLD (only taking measures before first LLD use), or 3) measurements of samples who only have records while taking LLD (adjusting measures for LLD).

We found that 127,903 samples had taken a LLD at some point of the following major categories: statins, fibrates, bile acid sequestrants, niacin, and cholesterol absorption inhibitors. We accounted for this by using LLD correction factors^35–38^ (**Supplementary Table 7**) and applied the correction factor with the largest effect (if multiple LLD were being taken) to the lipid measurement.

After adjusting for LLD, we calculated the median value of measurements per sample, resulting in 152,294 LDL-C median measurements.

### Genetic Variant Annotation

We annotated all genetic variants using Ensembl Variant Effect Predictor (VEP)^39^, LOFTEE^40^ and UTRannotator^41^. Where possible, we assigned each variant to one of three classifications: coding, proximal-regulatory or intergenic-regulatory. A variant was classified as coding if it had a predicted impact on the coding sequence of any transcript; proximal-regulatory if the variant lay within a 5 kb window of the UTRs of a transcript, and was not already a coding variant in any transcript, and finally regulatory if was not coding, but we did not assume proximality to coding transcript. We additionally tested variants in sliding windows of size 2000 base pairs, regardless of the number of variants in each window, with coding variants excluded to minimise hypothesis-testing overlap.

We then assigned each variant to groupings, which we refer to as masks, according to their predicted consequence and location. We used five published variant scores to group variants by consequence:

1. **Genomic Evolutionary Rate Profiling (GERP)** The GERP score is a measure of conservation at the variant level^42^. We classified a variant as highly conserved if it had a GERP score > 2.
2. **phastCon score** phastCon is a window-based measure of conservation across species^43^: either strictly mammalian (phastCon 30), or for all species (phast_100). We tested non-coding genome windows, i.e. excluding any window containing an exon, that had a phastCon score in the 99^th^ percentile.
3. **Gnomad 1 kb windows with Constraint Score** Constraint was calculated in windows of size 1 kb^25^ based on the local mutability and observed mutation rate of each window.
4. **SpliceAI score** The SpliceAI score^44^ is a measure of how well predicted each variant within a pre-mRNA region is of being a splice donor/acceptor, or neither. A variant was classified as a splice site with high confidence if it had an AI > 50.
5. **Combined Annotation Dependent Deletion score (CADD)** The CADD score predicts how deleterious a variant is likely to be^45^. We applied the CADD score only to coding variants and considered loss-of-function variants only if tagged as high confidence by VEP. Missense variants with CADD > 25 were segregated for testing in a separate mask.
6. ***JARVIS* Score** The JARVIS score^46^ was derived to better prioritise non-coding genetic variation for association study, based on a machine learning model derived from measures of constraint.

Each genome mask consisted of variants with different consequences, based on their location, one of the above scores and/or predicted coding consequences. For example, for a variant to be classified as missense CADD > 25, it must change a codon of an exon of a gene transcript and be predicted to be highly deleterious.

In **Supplementary Table 8** we present the full list of consequences assigned to each mask and classification.

### Association Analyses

We performed both single variant and aggregate tests genome-wide using REGENIE ^15^ (v4.1). All association analyses in UKB were corrected for age, sex, age squared, whole-genome sequencing batch, UKB recruitment centre and the first forty genetic principal components. All association analyses in AoU were corrected for age, sex, age squared, whole-genome sequencing batch, sample type (blood or saliva) and the first sixteen genetic principal components. Association analyses were also conditioned on all lead genetic variants reported in the most recent meta-analysis for LDL-C^1^, provided they were found within in the relevant whole-genome sequencing dataset. The LDL-C phenotype was rank inverse normalised at runtime in each stratum separately using REGENIEs --apply-rint option.

To increase the phenotypic variance explained by REGENIE’s leave-one-chromosome-out (LOCO) step-1 predictions we additionally adjusted for a leave-one-chromosome-out PGS generated from the most recent GLGC consortia common variant GWAS of LDL-C. This increases power in REGENIE step-2 association testing ^47^, particularly in smaller strata.

### Single Variant Association Testing and Meta-Analysis

To identify single variants (indels and single-nucleotide variants) associated with LDL-C we first performed association testing stratified by biobank-ancestry combination (UKB-AFR, UKB-EUR, UKB-SAS, AOU-AFR, AOU-AMR, AOU-EAS, AOU-EUR, AOU-MID & AOU-SAS). Within each stratum we tested all variants with MAC ≥ 11 using REGENIE (v4.1).

We then performed a fixed-effect meta-analysis of single variants with MAC ≥ 11 in any stratum using GWAMA^17^. GWAMA calculates the meta-frequency of a variant as the weighted average effect-allele frequency, in strata where the variant was tested, and this is the frequency we report in our results.

This initial single variant meta-analysis, which we used as the starting point for federated conditioning, did not adjust for variants from previous GWAS. We also ran a separate meta-analysis adjusting for previous GWAS variants to determine which of our results highlighted new loci.

### Aggregate Association Testing and Meta-Analysis

We next performed aggregate burden association testing for variants with two separate frequency thresholds: 1) MAF < 0.1% and 2) MAC ≤ 10, within each stratum, using REGENIE (v4.1) and our genetic annotations described above. Rare-variant aggregate burden effects were then meta-analysed using GWAMA.

To generate ACAT summary statistics we combined the meta-analysed test statistics from MAC ≤ 10 burden associations and all single variants with MAC > 10 and meta-MAF < 0.1%, using GCTA’s ACAT implementation^18^. We chose not to meta-analyse per-stratum ACAT p-values due to concerns about over-sensitivity.

This initial aggregate meta-analysis, which we used as the starting point for federated conditioning, adjusted for variants from previous GWAS. We also ran a separate meta-analysis for our independent aggregates, adjusting for previous GWAS variants as well as independent single variants identified by federated conditioning, to determine which of our aggregate results highlighted associations not picked up by our single variant analysis.

### Establishing single variant independence

To define a list of independent single variants, we applied a federated cross-biobank forward selection conditioning approach. We first selected the most strongly associated variant per chromosome. For each chromosome, within each stratum, we then repeated the association analyses for all other variants with *P*_*meta*_ < 1e-7 on that chromosome, conditioned on the selected variant (using REGENIE’s --condition-list option). We then meta-analysed the conditional results and selected the next most strongly associated variant that was not collinear with previously selected variant(s), per chromosome, using the variance inflation factor to determine collinearity with an r^2^ threshold of 0.9, similarly to the GCTA-CoJo forward step^19^. This process is repeated, always conditioning on the full set of selected variants, until no single variant associations have a meta-analysis p-value below the threshold for testing. Finally, we performed, in each stratum, a joint analysis of all the genome-wide significant selected variants using GCTA-CoJo and meta-analysed the results using GWAMA.

### Establishing aggregate independence

To define a list of independent aggregates, we applied a similar approach to that for single variants. We first selected the most strongly associated aggregate, from the union of burden and ACAT test results, per chromosome. For each chromosome, within each stratum, we then repeated the single variant (MAC > 11; MAF < 0.1%) and aggregate (BURDEN; MAC ≤ 10; MAF < 0.1%) analyses for all other aggregates with *P*_*meta*_ < 1e-7 on that chromosome, conditioned on all variants contributing to the selected aggregate (using REGENIE’s --condition-list option). We then meta-analysed the conditional results and selected the next most strongly associated aggregate, per chromosome. This process is repeated, always conditioning on the full set of selected aggregates, until no aggregate associations have a meta-analysis p-value below the threshold for testing. At each iteration, we dropped stratum-chromosome combinations where the number of variants being conditioned on exceeded half the stratum’s sample size.

### LDLR Locus Simulations

We selected the aggregate ‘LDLR (ENST00000557933) Missense (CADD > 25)’, comprised of 298 ultra-rare variants, as the causal aggregate. We then simulated 5 causal variant architectures with varying distributions of effect size (E): uniform (p(E = 1) = 1); sparse (p(E = 1) = 0.2, p(E = 0) = 0.8); bidirectional (p(E = 1) = p(E = -1) = 0.5), sparse bidirectional (p(E = 1) = p(E = -1) = 0.1, p(E = 0) = 0.8); normal (p(E) ∼ N(0, 1)). Using the same effect sizes in all strata, we used PLINK^33^ to calculate an effect size sum for each individual, before adding noise from normal distributions with standard deviations of 0.5, 1.0, 2.0 and 4.0 for a total of 20 phenotypes. We then performed association testing and federated conditioning on chromosome 19 following the methods in our primary analysis.

## Notes

https://github.com/ExeterGenetics/federated-conditioning-ldl-c-2026

### Competing Interest Statement

The authors have declared no competing interest.

## References

1. Graham, S. E. et al. The power of genetic diversity in genome-wide association studies of lipids. Nature 2021 600:7890 600, 675–679 (2021).

2. Karczewski, K. J. et al. Pan-UK Biobank genome-wide association analyses enhance discovery and resolution of ancestry-enriched effects. Nat. Genet. 1–10 (2025) doi:10.1038/S41588-025-02335-7;TECHMETA.

3. Wojcik, G. L. et al. Genetic analyses of diverse populations improves discovery for complex traits. Nature 570, 514–518 (2019).

4. Hawkes, G. et al. Whole-genome sequencing in 333,100 individuals reveals rare noncoding single variant and aggregate associations with height. Nat. Commun. 15, 8549 (2024).

5. Hawkes, G. et al. Whole-genome sequencing analysis identifies rare, large-effect noncoding variants and regulatory regions associated with circulating protein levels. Nature Genetics 2025 57:3 57, 626–634 (2025).

6. Ribeiro, D. M., Hofmeister, R. J., Rubinacci, S. & Delaneau, O. Noncoding rare variant associations with blood traits in 166,740 UK Biobank genomes. Nature Genetics 2025 57:9 57, 2146–2155 (2025).

7. Yengo, L. et al. A saturated map of common genetic variants associated with human height. Nature 610, (2022).

8. Joseph, T. A. et al. Computationally efficient meta-analysis of gene-based tests using summary statistics in large-scale genetic studies. Nature Genetics 2025 1–8 (2025) doi:10.1038/s41588-025-02390-0.

9. Li, X. et al. Powerful, scalable and resource-efficient meta-analysis of rare variant associations in large whole genome sequencing studies. Nature Genetics 2022 55:1 55, 154–164 (2022).

10. Turkmen, A. & Lin, S. Are rare variants really independent? Genet. Epidemiol. 41, 363–371 (2017).

11. Bycroft, C. et al. The UK Biobank resource with deep phenotyping and genomic data. Nature 562, 203–209 (2018).

12. Whole-genome sequencing of 490,640 UK Biobank participants. Nature 2025 645:8081 645, 692–701 (2025).

13. Bick, A. G. et al. Genomic data in the All of Us Research Program. Nature 2024 627:8003 627, 340–346 (2024).

14. The “All of Us” Research Program. N. Engl. J. Med. 381, 668–676 (2019).

15. Mbatchou, J. et al. Computationally efficient whole-genome regression for quantitative and binary traits. Nat. Genet. 53, 1097–1103 (2021).

16. Liu, Y. et al. ACAT: A Fast and Powerful p Value Combination Method for Rare-Variant Analysis in Sequencing Studies. Am. J. Hum. Genet. 104, 410–421 (2019).

17. Mägi, R. & Morris, A. P. GWAMA: Software for genome-wide association meta-analysis. BMC Bioinformatics 11, (2010).

18. Yang, J., Lee, S. H., Goddard, M. E. & Visscher, P. M. GCTA: A tool for genome-wide complex trait analysis. Am. J. Hum. Genet. 88, 76–82 (2011).

19. Yang, J. et al. Conditional and joint multiple-SNP analysis of GWAS summary statistics identifies additional variants influencing complex traits. Nat. Genet. 44, 369–375 (2012).

20. Harrell, F. E. Regression Modeling Strategies. https://doi.org/10.1007/978-3-319-19425-7 (2015) xdoi:10.1007/978-3-319-19425-7.

21. Vilariño-Güell, C. et al. DNAJC13 mutations in Parkinson disease. Hum. Mol. Genet. 23, 1794–1801 (2014).

22. Yoshida, S. et al. Parkinson’s disease-linked DNAJC13 mutation aggravates alphasynuclein-induced neurotoxicity through perturbation of endosomal trafficking. Hum. Mol. Genet. 27, 823–836 (2018).

23. Bjornsson, E. et al. Lifelong Reduction in LDL (Low-Density Lipoprotein) Cholesterol due to a Gain-of-Function Mutation in LDLR. Circ. Genom. Precis. Med. 14, E003029 (2021).

24. Gretarsdottir, S. et al. A Splice Region Variant in LDLR Lowers Non-high Density Lipoprotein Cholesterol and Protects against Coronary Artery Disease. PLoS Genet. 11, (2015).

25. Chen, S. et al. A genomic mutational constraint map using variation in 76,156 human genomes. Nature 2023 625:7993 625, 92–100 (2023).

26. Lonsdale, J. et al. The Genotype-Tissue Expression (GTEx) project. Nature Genetics 2013 45:6 45, 580–585 (2013).

27. GTEx Portal. https://www.gtexportal.org/home/gene/C8B.

28. Barton, A. R., Sherman, M. A., Mukamel, R. E. & Loh, P. R. Whole-exome imputation within UK Biobank powers rare coding variant association and fine-mapping analyses. Nat. Genet. 53, 1260–1269 (2021).

29. Danecek, P. et al. Twelve years of SAMtools and BCFtools. Gigascience 10, 1–4 (2021).

30. Poterba, T. et al. The scalable variant call representation: enabling genetic analysis beyond one million genomes. Bioinformatics 41, (2024).

31. All of Us Genomic Quality Report – User Support. https://support.researchallofus.org/hc/en-us/articles/29390274413716-All-of-Us-Genomic-Quality-Report.

32. gatk/scripts/variantstore/docs/release_notes/VETS_Release.pdfatah_var_store ·broadinstitute/gatk. https://github.com/broadinstitute/gatk/blob/ah_var_store/scripts/variantstore/docs/release_notes/VETS_Release.pdf.

33. Chang, C. C. et al. Second-generation PLINK: Rising to the challenge of larger and richer datasets. Gigascience 4, (2015).

34. Chen, H. H., Petty, L. E., Gamazon, E. R., Wells, Q. S. & Below, J. E. Optimizing Genetic Analyses of Serum Lipids in Longitudinal Data. Circ. Res. 127, 1337–1339 (2020).

35. Wu, J. et al. An investigation of the effects of lipid-lowering medications: genome-wide linkage analysis of lipids in the HyperGEN study. BMC Genet. 8, (2007).

36. Third Report of the National Cholesterol Education Program (NCEP) Expert Panel on Detection, Evaluation, and Treatment of High Blood Cholesterol in Adults (Adult Treatment Panel III) Final Report. Circulation 106, 3143–3421 (2002).

37. Sweeney, M. E. & Johnson, R. R. Ezetimibe: an update on the mechanism of action, pharmacokinetics and recent clinical trials. Expert Opin. Drug Metab. Toxicol. 3, 441–450 (2007).

38. Hung-Hsin Chen, B. & Below Nancy J Cox Kari E North Quinn S Wells PharmD Todd L Edwards, J.E. Comprehensively Investigating the Genetic Architecture of Serum Lipids: From Identity by Descent Mapping, Transcriptome-Wide Association Study to Phenome-Wide Association Study. Preprint at http://hdl.handle.net/1803/16051 (2020).

39. McLaren, W. et al. The Ensembl Variant Effect Predictor. Genome Biol. 17, 1–14 (2016).

40. Karczewski, K. J. et al. The mutational constraint spectrum quantified from variation in 141,456 humans. Nature 581, 434–443 (2020).

41. Zhang, X., Wakeling, M., Ware, J. & Whiffin, N. Annotating high-impact 5′untranslated region variants with the UTRannotator. Bioinformatics 37, 1171–1173 (2021).

42. Davydov, E. V. et al. Identifying a high fraction of the human genome to be under selective constraint using GERP++. PLoS Comput. Biol. 6, (2010).

43. Siepel, A. et al. Evolutionarily conserved elements in vertebrate, insect, worm, and yeast genomes. Genome Res. 15, 1034–1050 (2005).

44. Jaganathan, K. et al. Predicting Splicing from Primary Sequence with Deep Learning. Cell 176, 535-548.e24 (2019).

45. Rentzsch, P., Witten, D., Cooper, G. M., Shendure, J. & Kircher, M. CADD: Predicting the deleteriousness of variants throughout the human genome. Nucleic Acids Res. 47, D886– D894 (2019).

46. Vitsios, D., Dhindsa, R. S., Middleton, L., Gussow, A. B. & Petrovski, S. Prioritizing noncoding regions based on human genomic constraint and sequence context with deep learning. Nat. Commun. 12, 1–14 (2021).

47. Campos, A. I. et al. Boosting the power of genome-wide association studies within and across ancestries by using polygenic scores. Nat. Genet. 55, 1769–1776 (2023).

