## Supplementary Figures 1-4 for "Federated cross-biobank conditional analysis identifies LDL-C lowering effects of *DNAJC13* haploinsufficiency and *LDLR* regulation"

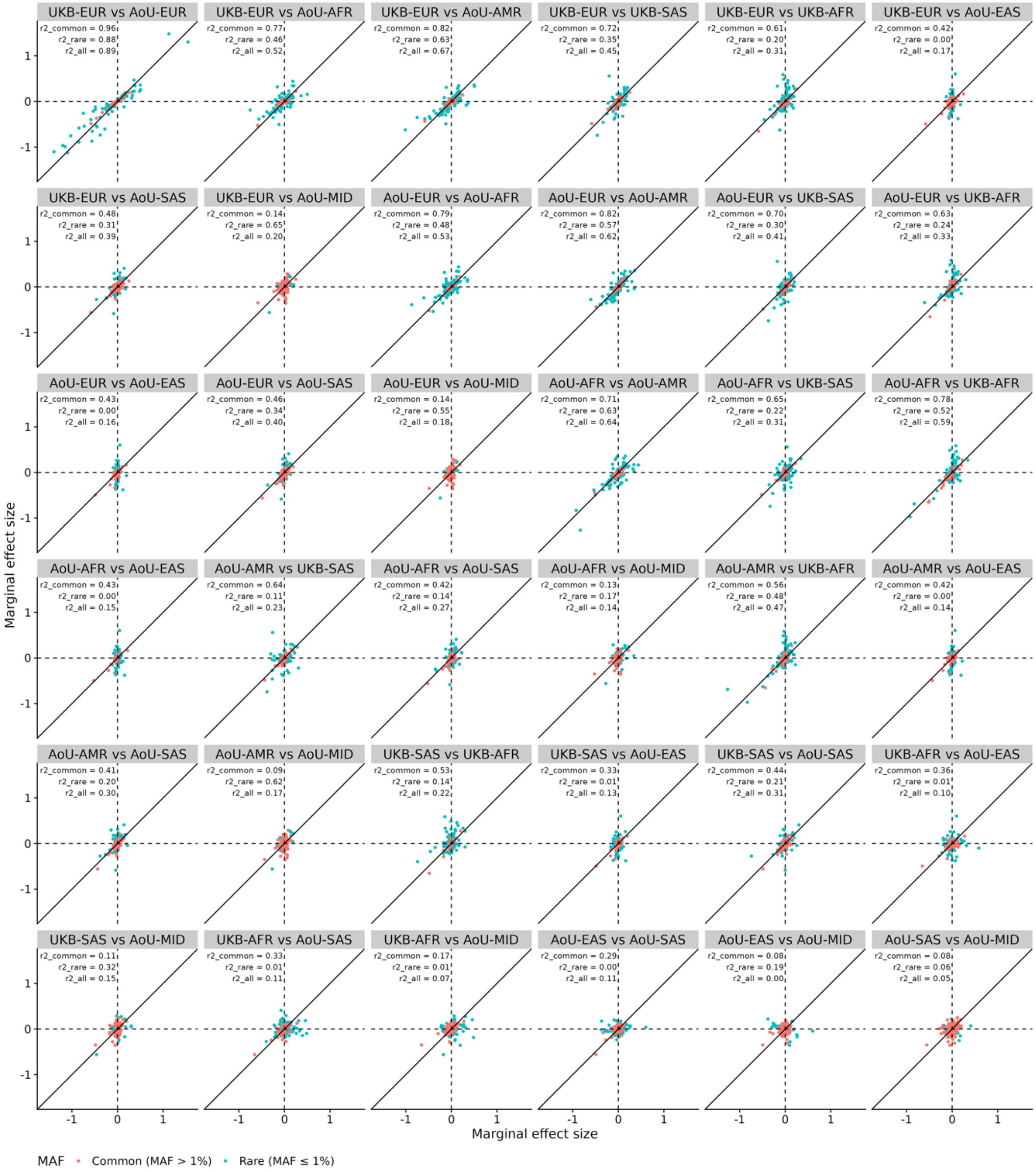


**Supplementary Figure 1: Single variant association effect-size concordance for LDL-C.** Scatter plots showing effect size concordance across biobanks and ancestries for independent single-variant associations. Each panel compares effect sizes for the strata in the title: the effect size of the first of two strata is plotted on the x-axis. Red points represent common variants, and blue points represent rare variants (<1% MAF in either stratum).

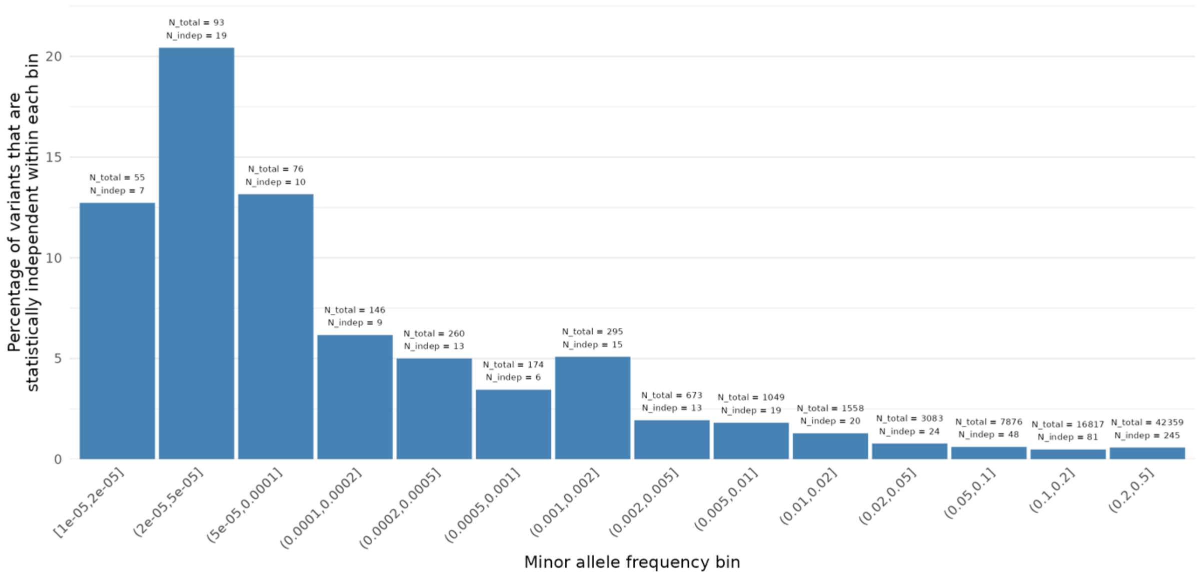


**Supplementary Figure 2: Rarer variants are more likely to be statistically independent than common variants, with up to 21% of the rarest variants being independent.** Bar plot showing the percentage of variants significant in the unconditional meta-analysis which were determined as independently significant by meta-conditioning, across distinct minor allele frequency bins. Labels show the number variants that were significant before (N_total) and after (N_indep) meta-conditioning.

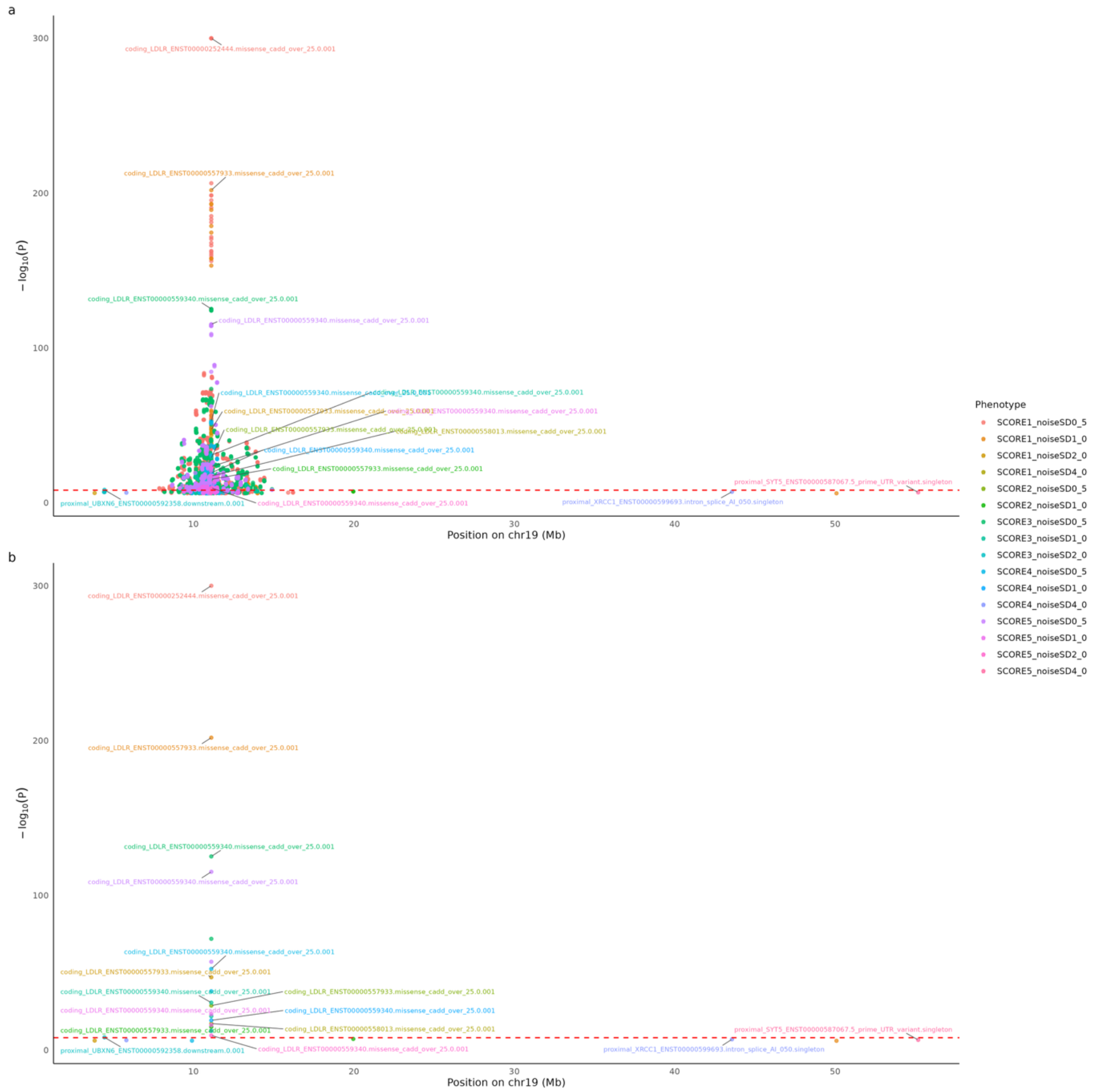


**Supplementary Figure 3: Meta-conditioning of aggregates supresses spurious associations.** Manhattan plots of aggregate associations a) before conditioning and b) after conditioning. The true causal aggregate is ‘LDLR (ENST00000557933) Missense (CADD > 25)’, comprised of 298 ultra-rare variants. The 20 colours differentiate the 20 simulated phenotypes, including 5 causal variant architectures (SCORES) with varying distributions of effect size (E): SCORE1 uniform (p(E = 1) = 1); SCORE2 sparse (p(E = 1) = 0.2, p(E = 0) = 0.8); SCORE3 bidirectional (p(E = 1) = p(E = -1) = 0.5); SCORE4 sparse bidirectional (p(E = 1) = p(E = -1) = 0.1, p(E = 0) = 0.8); SCORE5 normal (p(E) ~ N(0, 1)), as well as 4 levels of normally distributed noise, with standard deviations 0.5, 1.0, 2.0 and 4.0, for a total of 20 phenotypes.

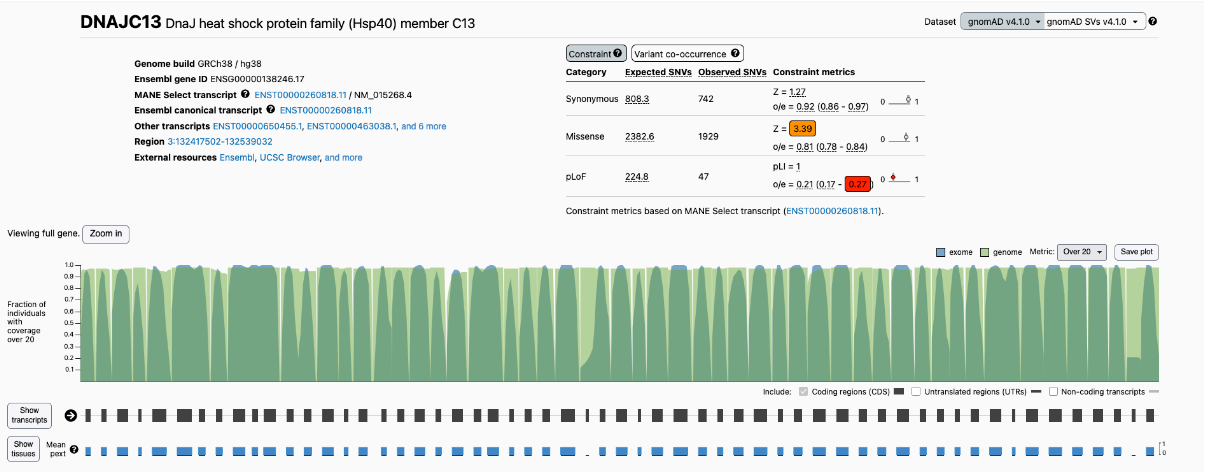


**Supplementary Figure 4: Sequencing coverage for DNAJC13.** GNOMAD coverage (v4.1.0, accessed 01/12/2025) over the exonic regions of *DNAJC13* for whole-genome sequencing (green) and whole-exome sequencing (blue).
